# CLEESE: An open-source audio-transformation toolbox for data-driven experiments in speech and music cognition

**DOI:** 10.1101/436477

**Authors:** Juan José Burred, Emmanuel Ponsot, Louise Goupil, Marco Liuni, JJ Aucouturier

## Abstract

Over the past few years, the field of visual social cognition and face processing has been dramatically impacted by a series of data-driven studies employing computer-graphics tools to synthesize arbitrary meaningful facial expressions. In the auditory modality, reverse correlation is traditionally used to characterize sensory processing at the level of spectral or spectro-temporal stimulus properties, but not higher-level cognitive processing of e.g. words, sentences or music, by lack of tools able to manipulate the stimulus dimensions that are relevant for these processes. Here, we present an open-source audio-transformation toolbox, called CLEESE, able to systematically randomize the prosody/melody of existing speech and music recordings. CLEESE works by cutting recordings in small successive time segments (e.g. every successive 100 milliseconds in a spoken utterance), and applying a random parametric transformation of each segment’s pitch, duration or amplitude, using a new Python-language implementation of the phase-vocoder digital audio technique. We present here two applications of the tool to generate stimuli for studying intonation processing of interrogative vs declarative speech, and rhythm processing of sung melodies.

## Introduction

The field of high-level visual and auditory research is concerned with the sensory and cognitive processes involved in the recognition of objects or words, faces or speakers and, increasingly, of social expressions of emotions or attitudes in faces, speech and music. In traditional psychological methodology, the signal features that drive judgments (e.g. facial metrics such as width-to-height ratio, acoustical features such as mean pitch) are posited by the experimenter before being controlled or tested experimentally, which may create a variety of confirmation biases or experimental demands. For instance, stimuli constructed to display western facial expressions of happiness or sadness may well be recognized as such by non-western observers [1], but yet may not be the way these emotions are spontaneously produced, or internally represented, in such cultures [2].

Similarly in auditory cognition, musical stimuli recorded by experts pressed to express emotions in music may do so by mimicking expressive cues used in speech, but these cues may not exhaust the many other ways in which arbitrary music can express emotions [3]. For all these reasons, in recent years, a series of powerful data-driven paradigms (built on techniques such as reverse-correlation, classification image or bubbles; see [4] for a review) were introduced in the field of visual cognition to discover relevant signal features empirically, by analyzing participant responses to large sets of systematically-varied stimuli [5].

**Fig 1.**
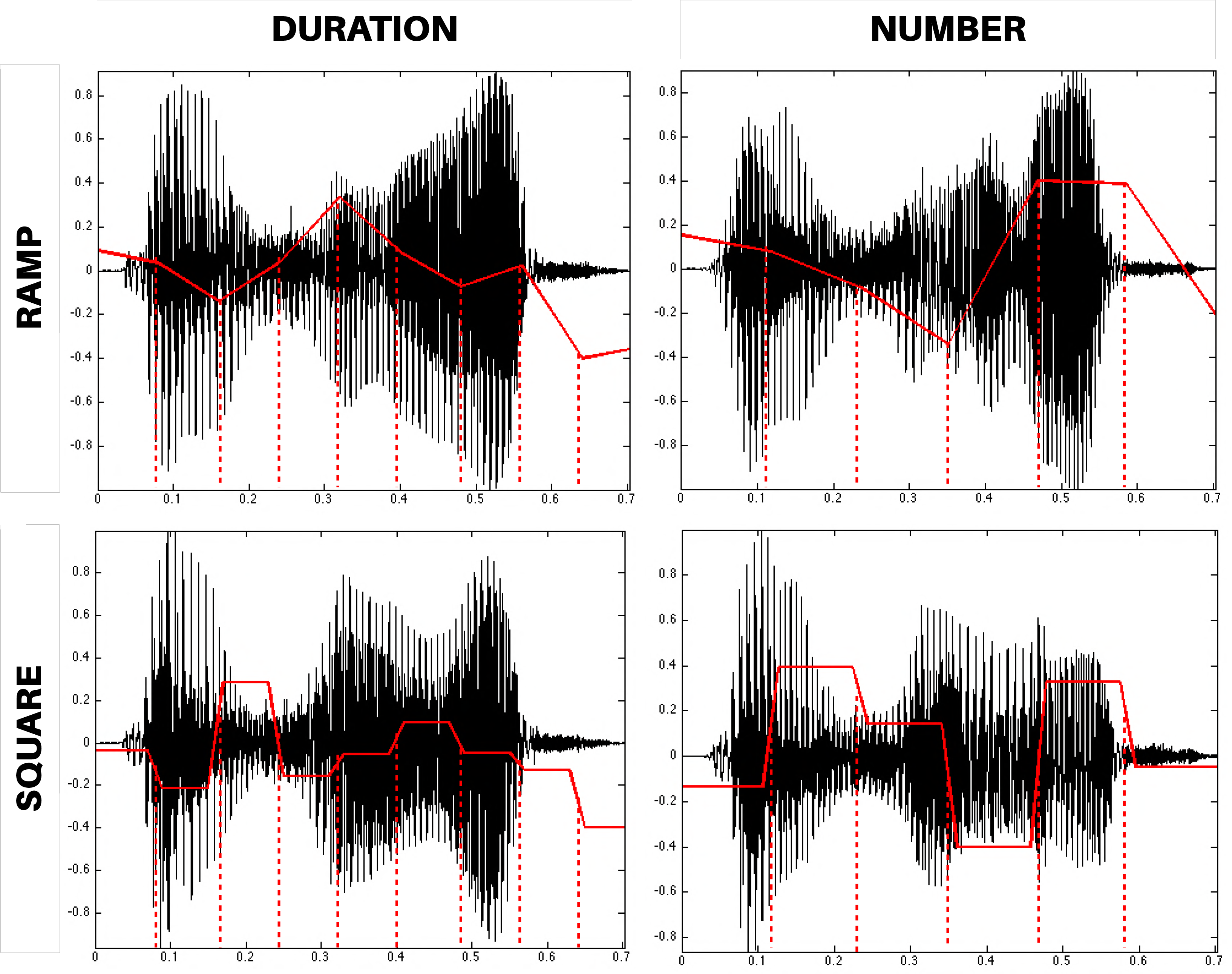
Low- and high-level subspace noise for the reverse-correlation of visual and audio stimuli. Reverse-correlation paradigms aim to isolate the subspace of feature dimensions that maximizes participant responses, and as such, need to search a stimulus generative space e.g. of all possible images or sounds relevant for a task. In a majority of studies, noise masks operate on low-level, frequency-based representations of the signal, e.g. at different scales and orientations for image stimuli (top-left) or different frequencies of an auditory spectrogram (top-right). More recent models in the vision modality are able to explore the subspace of facial expressions through the systematic manipulation of facial action units (bottom-left). The present work represents a conceptually-similar development for the auditory modality.

The reverse correlation technique was first introduced in neurophysiology to characterize neuronal receptive fields of biological systems with so-called “white noise analysis” [6–9]. In psychophysics, the technique was then adapted to characterize human sensory processes, taking behavioral choices (e.g., yes/no responses) instead of neuronal spikes as the systems’ output variables to study, e.g. in the auditory domain, detection of tones in noise [10] or loudness weighting in tones and noise ([11]; see [4] for a review of similar applications in vision). In the visual domain, these techniques have been extended in recent years to address not only low-level sensory processes, but higher-level cognitive mechanisms in humans: facial recognition [12], emotional expressions [2, 13], social traits [14], as well as their associated individual and cultural variations ([15]; for a review, see [5]). In speech, even more recently, reverse correlation and the associated “bubbles” technique were used to study spectro-temporal regions underlying speech intelligibility [16, 17] or phoneme discrimination in noise [18, 19] and, in music, timbre recognition of musical instruments [20, 21].

All of these techniques aim to isolate the subspace of feature dimensions that maximizes participant responses, and as such, need to search the stimulus generative space e.g. of all possible images or sounds relevant for a given task. A typical way to define and search such a space is to consider a single target stimulus (e.g. a neutral face, or the recording of a spoken phoneme), apply a great number of noise masks that modify the low- or high-level physical properties of that target, and then regress the (random) physical properties of the masks on participant judgements. Techniques differ in how such noise masks are generated, and on what stimulus subspace they operate (Figure 1). At the lowest possible level, early proposals have applied simple pixel-level noise masks [13], and sometimes even no stimulus at all, akin to white noise “static” on a TV screen in which participants were forced to confabulate the presence of a visual target [22] or a vowel sound [19]. More consistently, noise masks generally operate on low-level, frequency-based representations of the signal, e.g. at different scales and orientations for image stimuli [12, 14], different frequencies of an auditory spectrogram [17, 23] or different rates and scales of a modulation spectrum (MPS) [16, 21].

While low-level subspace noise has the advantage of providing a physical description of the stimuli driving participant responses, it is often a suboptimal search space for high-level cognitive tasks. First, all stimulus features that are driving participant judgements may not be efficiently encoded in low-level representations: the auditory modulation spectrum, for instance, is sparse for a sound’s harmonic regularities, coded for by a single MPS pixel, but not for features localized in time, such as attack time or transitions between phonemes [23]. As a consequence, local mask fluctuations will create continuous variations for the former, but not the latter features which will be difficult to regress on. Second, low-level variations in stimuli typically create highly distorted faces or sounds (Figure 1-top), for which one may question the ecological relevance of participant judgements. Finally, and perhaps most critically, many of the most expressive, cognitively-meaningful features of a face or voice signal are coordinated action-units (e.g. a contraction of a facial muscle, the rise of pitch at the end of a spoken utterance) that have distributed representations in low-level search spaces, and will never be consistently explored by a finite amount of random activations at that level.

Consequently, the face research community has recently developed a number of higher-level generative models able to synthesize facial expressions through the systematic manipulation of facial action units [24, 25]. These tools are based on morphological models that simulate the ‘amplitude vs time’ effect of individual muscles and then reconstruct realistic facial textures that account for the modified underlying morphology, as well as head pose and lighting. Searching such high-level subspaces with reverse-correlation is akin to searching the space of all *possible* facial expressions (Figure 1-bottom), while leaving out the myriad of other possible facial stimuli that are not directly interpretable as human-made expressions - a highly efficient strategy that has been applied to characterize subtle aspects of social and emotional face perception processes, such as cultural variations in emotional expressions [2], physical differences between different kinds of smiles [26] or in the way these features are processed in time [27]. In auditory research, however, similarly efficient data-driven strategies have not yet been common practice, by lack of tools able to manipulate the high-level stimulus dimensions that are relevant for similar judgement tasks when they apply on, e.g., voice or music.

Here, we present an open-source audio-transformation toolbox, called CLEESE (Combinatorial Expressive Speech Engine) ^1^, able to systematically randomize the prosody/melody of existing speech and music recordings. CLEESE works by cutting recordings in small successive time segments (e.g. every successive 100 milliseconds in a spoken utterance), and applying a random parametric transformation of each segment’s pitch, duration, amplitude or frequency content, using a new Python-language implementation of the phase-vocoder digital audio technique. Transformations made with CLEESE explore the space of speech intonation and expressive speech prosody by allowing to create random time-profiles of pitch (e.g. rising pitch at the end of an utterance, as in interrogative sentences [28]), of duration (e.g. word-final vowel lengthening, as used as a cue for word segmentation [29]) or amplitude (e.g. louder on prominent words or phonemes [30]). In music, the same transformations can be described as melodic, manipulating the pitch/tuning of successive notes in a sequence (e.g. ♩♭, ♩ or ♩♯), their duration (e.g. 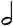, ♩ or ♪) or amplitude (e.g. *p*, *mf* or *f*). All transformations are parametric, thus allowing to generate thousands of random variants drawn e.g. from a gaussian distribution; and realistic (within appropriate parameter ranges), such that the resulting audio stimuli do not typically appear artificial/transformed, but rather plausible as ecological speech or music recordings.

In recent work, we have used CLEESE with a reverse-correlation paradigm to uncover what mental representations of pitch profiles underlie judgements of speaker dominance and trustworthiness in short utterances like the word ‘hello’ [31]. We recorded a single utterance of the word ‘hello’ by one male and one female speaker. CLEESE was first used to flatten the pitch of the recordings (by transforming it with a pitch profile that alters its original prosody to constant pitch), and then to generate random pitch variations, by manipulating the pitch over 6-time points on Gaussian distributions of SD = 70 cents clipped at ±2.2 SD. Pairs of these randomly-manipulated voices were then presented to observers who were asked, on each trial, to judge which of the two versions appeared most dominant/trustworthy. The participants’ mental representations were then computed by computing the mean pitch contour (a 6-point vector) of the voices they classified as dominant (resp. trustworthy) minus the mean pitch contour of the voices classified as non-dominant (resp. non-trustworthy).

In this article, we describe the functionality of the CLEESE toolbox, the algorithms that underlie it as well as how to deploy it in psychophysical reverse-correlation experiments such as those above. We then present two case-studies in which we use CLEESE to generate pitch-shifted speech stimuli to study the perception of interrogative vs declarative speech prosody, and time-stretched musical stimuli to study the rhythm processing of sung melodies.

## Functionality, algorithms, usage

### Functionality

CLEESE is an open-source Python toolbox^2^ used to create random or fixed pitch, time and amplitude transformations on any input sound^3^. The transformations can be both static or time-varying. Besides its purpose of generating many stimuli for reverse-correlation experiments, the toolbox can also be used for producing individual, user-determined transformations.

CLEESE operates by generating a set of random breakpoint functions (BPFs) for each transformation, which are then passed to a spectral processing engine (based on the phase vocoder algorithm, see below) for the transformation to occur. BPFs determine how the sound transformations vary over the duration of the stimulus.

CLEESE can randomly generate BPFs of two types: *ramps*, where the corresponding sound parameter is interpolated linearly between breakpoints (Figure 2-top) and *square*, where the BPF is a square signal with sloped transitions (Figure 2-bottom). BPF segments can be defined either by forcing all of them to have a fixed duration (i.e., their number will depend on the whole sound’s duration, see Figure 2-left), or by forcing a fixed number of segments along the total sound duration (i.e., their duration will depend on the whole sound’s duration, see Figure 2-right). Alternatively, the user can pass custom breakpoint positions, which can be defined manually e.g. to correspond to each syllable or note in given a recording.

**Fig 2.**
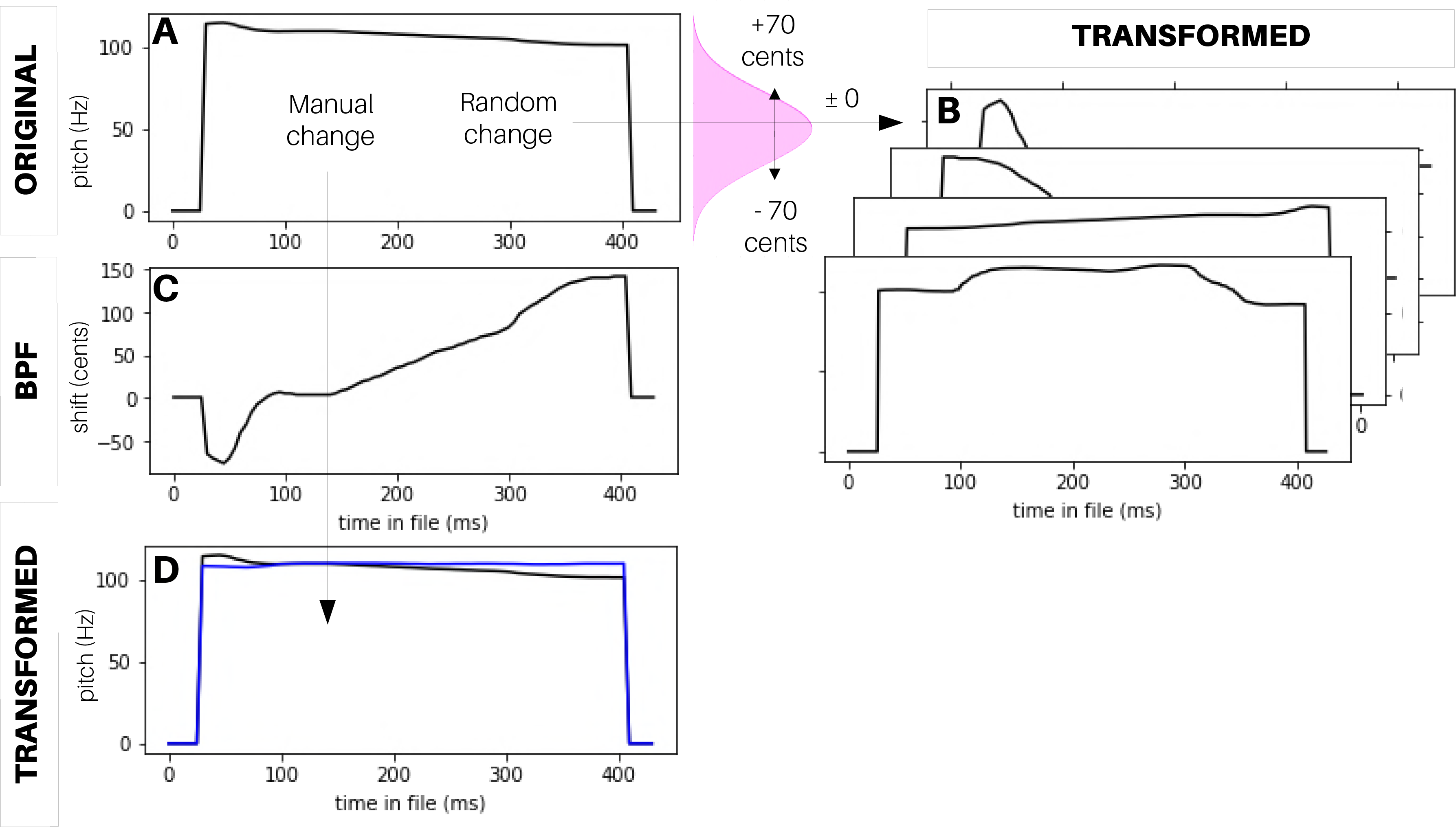
CLEESE operates by generating a set of random breakpoint functions (BPFs) which control the dynamically-changing parameters of the sound transformations. In *ramp* BPFs (top), transformation parameters are interpolated linearly between breakpoints. In *square* BPFs, they are constant in each segment. BPFs can be specified either in terms of fixed duration (left: 80ms segments) or fixed number of segments (right: 6 segments).

BPFs can be defined to transform sounds along three signal dimensions:

**Pitch:** The BPF is used to transpose up and down the pitch of each segment in the sound, while maintaining its amplitude and duration constant. Each breakpoint in the BPF corresponds to a pitch shift value *p* in *cents* or percents of a semitone, with 0 cents corresponding to no change with respect to original pitch.

**Time:** The BPF is used to stretch or compress the duration of each segment in the sound, while maintaining its amplitude and pitch constant. Each breakpoint corresponds to a time stretch factor *t* in ratio to original duration, with 0 < *t* < 1 corresponding to compression/faster speed, *t >* 1 stretched/slower speed, and *t* = 1 no change with respect to segment’s original duration.

**Amplitude:** The BPF is used to amplify or decrease the signal’s instantaneous amplitude in each segment, while maintaining its pitch and duration constant. Each breakpoint corresponds to a gain value *g* in dBs (thus, in base-10 logarithm ratio to original signal amplitude), with *g* = 0 corresponding to no change with respect to original amplitude.

The default mode is for CLEESE to generate *p*,*t* or *g* values at each breakpoint by sampling from a Gaussian distribution centered on *p* = 0, *t* = 1 or *g* = 0 (no change) and whose standard deviation (in cents, duration or amplitude ratio) allows to statistically control the intensity of the transformations. For instance, with a pitch SD of 100cents, CLEESE will assign random shift values at every breakpoint, 68% of which are within ± 1 semitone of the segment’s original pitch (Figure 3A-B). The transformations can be chained, e.g. manipulating pitch, then duration, then amplitude, all with separate transformation parameters. For each type of transformation, distributions can be truncated (at given multiples of SD) to avoid extreme transformation values which may be behaviorally unrealistic. Alternatively, transformation parameters at each breakpoint can be provided manually by the user. For instance, this allows to flatten the pitch contour of an original recording, by constructing a custom breakpoint function that passes through the pitch shift values needed to shift the contour to a constant pitch value (Figure 3C-D).

**Fig 3.**
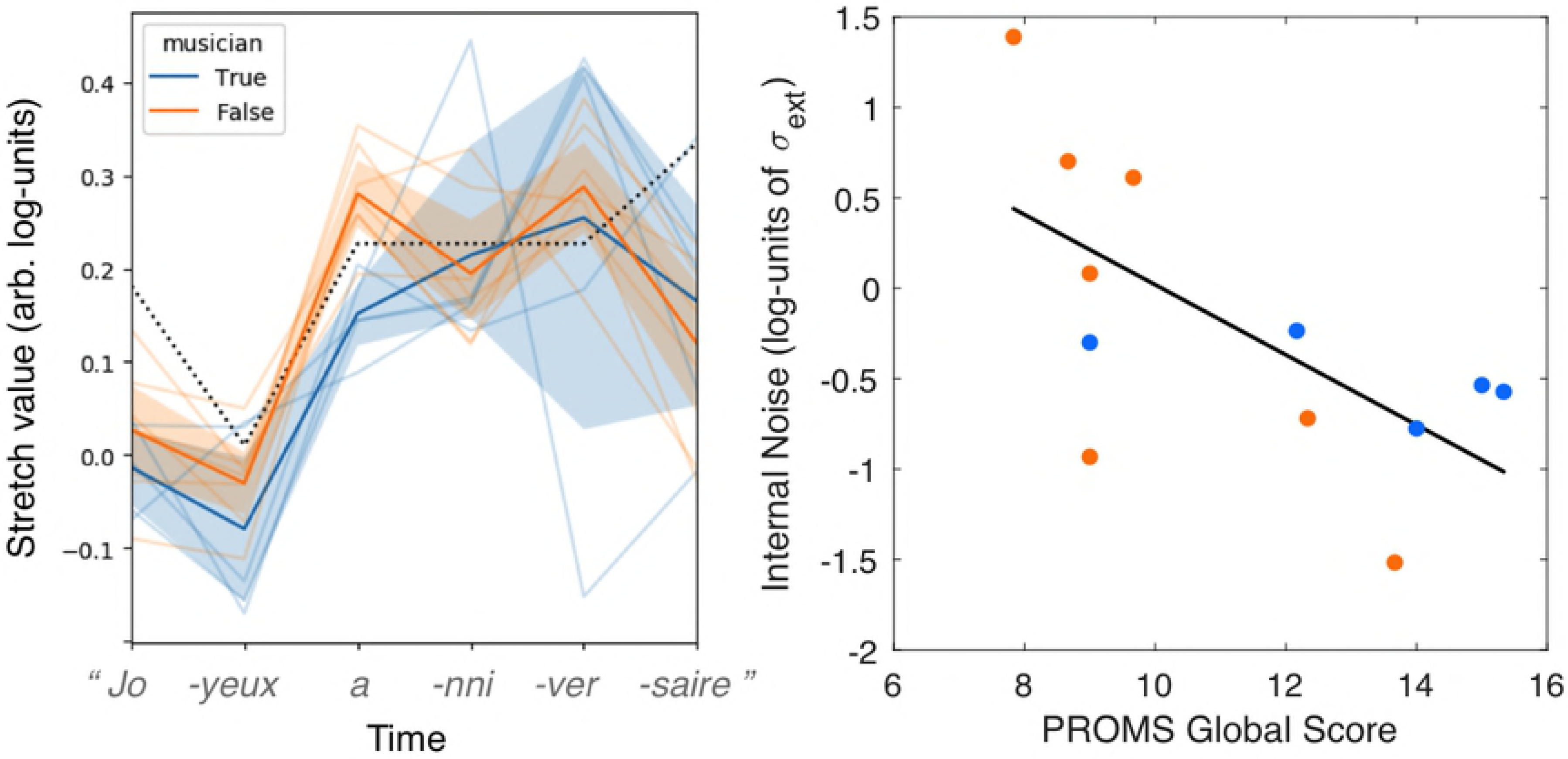
Examples of pitch manipulations created with CLEESE. A: pitch of an original male speech recording (a 400ms utterance of the word ‘hello’). B: Random pitch transformations of the original, where pitch shift values at each breakpoint are drawn from a Gaussian distribution. C: Manually-specified BPF created to flatten the pitch contour of the original recording. D: Resulting flat-pitch transformation, compared to original.

### Algorithm

The phase vocoder is a sound processing technique based on the short-term Fourier Transform (STFT, [33]). The STFT decomposes each successive parts (or *frames*) of an incoming audio signal into a set of coefficients that allow to perfectly reconstruct the original frame as a weighted sum of modulated sinusoidal components (eq.(1)). The phase vocoder algorithm operates on each frame’s STFT coefficients, modifying them either in their content (e.g. displacing frequency components to higher frequency positions to simulate a higher pitch) or their position in time (e.g. displacing frames to subsequent time points to simulate a slower sound). It then reconstructs the original signal from the manipulated frames with a variety of techniques meant to ensure the continuity (or phase coherence) of the resulting sinusoidal components [34, 35]. Depending on how individual frames are manipulated, the technique can be used e.g. to change a sound’s duration without affecting its pitch (a process known as *time stretching*), or to change a sound’s pitch without changing its duration (*pitch shifting*, see [36] for a review).

In more detail, the phase vocoder procedure considers a (digital) sound *x* as a real-valued discrete signal, and *h* a symmetric real-valued discrete signal composed of *N* samples, usually called *window*, that is used to cut the input sound into successive frames. In the analysis stage, frames are extracted with a time step (or *hop size*) *R*_*a*_, corresponding to successive time positions 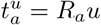, where *u* is the index of the *u*-th frame. The discrete STFT of *x* with window *h* is given by the discrete FFT of the time frames of *x* multiplied by *h*,

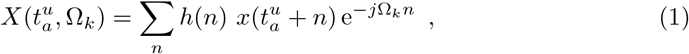

where 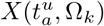 is the STFT coefficient corresponding to frequency 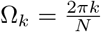 *k* = 0, …, *N* − 1 in the *u*-th frame of the signal. *X* is a two-dimensional complex-valued representation, which can be expressed in terms of real and imaginary parts or, equivalently, amplitude and phase. The amplitude of the coefficient at time index *u* and frequency bin *k* is given by 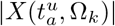, while its phase is 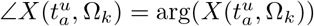.

A given transformation T(*X*) = *Y* can then be performed by altering the amplitudes and/or phases of selected coefficients, leading to a complex-valued representation *Y* of the same dimension as *X*. For instance, if one wishes to shift the frequency content of the sound by a fixed frequency *p*, then one can take 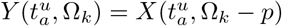 (i.e. the *k*-th frequency component of the *u*-th input frame is copied at the *k* + *p* frequency position in the *u*-th output frame). Alternatively, if we wish to temporally stretch/compress the sound by a given factor *t*, then we take 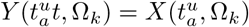 (i.e. the *k*-th frequency component of the *u*-th input frame is copied at the *k*-th frequency component of the *t* × *u*-th output frame, i.e. at a later time position if *t* > 1 or earlier position if *t* < 1). Finally, amplitude manipulations by a time-frequency mask *G* composed of scalar gain factors *g*_*u,k*_ can be generated by taking 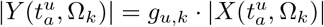 for the amplitudes, and leaving phases unchanged^4^.

The transformed coefficients in *Y* are then used to synthesize a new sound *y*, using the inverse procedure to eq.(1). As done for analysis, let the scalar *R*_*s*_ be the synthesis hop size, and 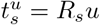 the position of the *u*-th output frame. Then, using the same window function *h*, the output signal is given by

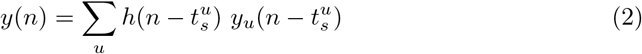

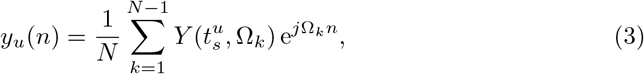

When STFT frames are time-shifted as part of the phase-vocoder transformation, the position of the *u*-th output frame is different from 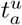 and the phases of the STFT coefficients have to be adapted to ensure the continuity of the reconstructed sinusoidal components and the perceptual preservation of the original sound’s timbre properties. Phase vocoder implementations offer several methods to this end [36]. In one classic procedure (*horizontal phase synchronization*), phases are adjusted independently in all frequency positions, with phases at position 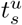 extrapolated from phases at position 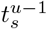 [34]. In an improved procedure (*vertical phase synchronization*, or *phase-locking*), each frame is first analysed to identify prominent peaks of amplitude along the frequency axis, and phases are only extrapolated in the peak frequency positions; phases of the frequency positions around each peak are locked to the phase of the peak frequency position [35]. It is this second procedure that is implemented in CLEESE.

### Usage

CLEESE is implemented as module for the Python (versions 2 and 3) programming language, and distributed under an open-source MIT License. It provides its own implementation of the phase vocoder algorithm, with its only dependencies being the `numpy` and `scipy` libraries.

In its default ‘batch’ mode, CLEESE generates many random modifications from a single input sound file, called the base sound. It can be launched as follows: 

~~~
import cleese
inputFile = ‘path_to_input_sound.wav’
configFile = ‘path_to_config_file.py’
cleese.process(soundData=inputFile,configFile=configFile)
~~~

where `inputFile` is the path to the base sound (a mono sound in WAV format) and `configFile` is the path to a user-generated configuration script containing generation parameters for all transformations. For each run in batch mode, the toolbox generates an arbitrary number of random BPFs for each transformation, applies them to the base sound, and saves the resulting files and their parameters.

The main parameters of the configuration file include how many files should be generated, where the transformed files should be saved, and what transformation (or combinations thereof) should be applied. For instance, the following configuration file generates 10 audio files which result from a random stretch, followed by a random pitch transformation of the base sound.

~~~
*# main parameters*
main_pars = {
        ‘numFiles’: 10, *# number of output files to generate*
        ‘outPath’: ‘/path_to_output_folder/’, *# output root folder*
        ‘chain’: True, *# apply transformations in series (True) # or parallel (False)*
        ‘transf’: [’stretch’,’pitch’] *# modifications to apply*
}
~~~

In addition, the configuration file includes parameters that specify how BPFs should be generated for each transformation, including the number or duration of BPF windows, their type (ramp or square) and the standard deviation of the gaussian distribution used to sample breakpoints. As an example, the following parameters correspond to pitch BPFs consisting of 6 ramp windows, each with a normally distributed pitch shift between −300 and 300 cents.

~~~
*# pitch transformation parameters*
pitch_pars = {
        ‘winLen’: 0.11, *# BPF window (sec)
        # if 0 : static transformation*
        ‘numWin’: 6, *# number of BPF windows.
        # if 0 : static transformation*
        ‘winUnit’: ‘n’, *# ‘s’: force winLen in seconds,
        # ‘n’: force number of windows*
        ‘std’: 300, *# standard deviation for each breakpoint*
        ‘trunc’: 1, *# truncate distribution (factor of std)*
        ‘BPFtype’: ‘ramp’, *# type of breakpoint function:
        # ‘ramp’: linear interpolation between breakpoints
        # ‘square’: square BPF*
        ‘trTime’: 0.02 *# transition time for square BPF (sec)*
}
~~~

The CLEESE module is distributed with a Jupyter notebook tutorial, showing further examples of using the toolbox for sound manipulation. In addition, all experimental data and analysis scripts (also in the form of Jupyter notebooks) from the following two case-studies are made available as supplementary material.

## Case-study (1): speech intonation

To illustrate the use of CLEESE with speech stimuli and pitch shifting, we describe here a short reverse-correlation experiment about the perception of speech intonation. Speech intonation, and notably the temporal pattern of pitch in a given utterance, can be used to convey syntactic or sentence mode information (e.g. whether a sentence is interrogative or declarative), stress (e.g. on what word is the sentence’s focus), emotional expression (e.g. whether a speaker is happy or sad) or attitudinal content (e.g. whether a speaker is confident or doubtful) [37]. For instance, patterns of rising pitch are associated with social traits such as submissiveness, doubt or questioning, and falling pitch with dominance or assertiveness [38–40]. In addition, recent neurophysiological evidence suggests that intonation processing is rooted at early processing stages in the auditory cortex [41]. However, it has remained difficult to attest of the generality of such intonation patterns and of their causality in cognitive mechanisms. Even for information as seemingly simple as the question/statement contrast, which is conventionally associated with the “final Rise” intonation, empirical studies show that, while frequent, this pattern is not as simple nor common as usually believed [42]. For instance, in one analysis of a corpus of 216 questions, the most frequent tone for polar questions (e.g. “Is this a question?”) was a Fall [43]. In addition, in English, interrogative pitch contours do not consistently rise on the final part of the utterance, but rather after the first syllable of the content word [44] (e.g. “Is this a *question you’re asking* ? vs “Is this a question you’re *asking* ?”).

We give here a proof of concept of how to use CLEESE in a reverse-correlation experiment to uncover what exact pitch contour drives participants’ categorization of an utterance as interrogative or declarative. Data come from the first experiment presented in [31].

## Methods

### Stimuli

One male speaker recorded a 426ms utterance of the French word “vraiment” (“really”), which can be experienced either as a one-word statement or question. We used CLEESE to artificially manipulate the pitch contour of the recording. First, the original pitch contour (mean pitch = 105Hz) was artificially flattened to constant pitch, using the procedure shown in Figure 3C-D. Then, we added/subtracted a constant pitch gain (±20 cents, equating to ± 1 fifth of a semitone) to create the ‘high-’ or ‘low-pitch’ versions presented in each trial^5^. Finally, we added Gaussian “pitch noise” (i.e. pitch-shifting) to the contour by sampling pitch values at 6 successive time-points, using a normal distribution (SD = 70 cents; clipped at ± 2.2 SD), linearly interpolated between time-points, using the procedure shown in Figure 3A-B.

### Procedure

700 pairs of randomly-manipulated voices were presented to each of N=5 observers (male: 3, M=22.5yo), all native French speakers with self-reported normal hearing.

Participants listened to a pair of two randomly-modulated voices and were asked which of the two versions was most interrogative. Inter-stimulus interval in each trial was 500 ms, and inter-trial interval was 1s.

### Apparatus

The stimuli were mono sound files generated at sampling rate 44.1 kHz in 16-bit resolution by Matlab. They were presented diotically through headphones (Beyerdynamic DT 770 PRO; 80 ohms) at a comfortable sound level.

### Analysis

A first-order temporal kernel [45] (i.e., a 7-points vector) was computed for each participant, as the mean pitch contour of the voices classified as interrogative minus the mean pitch contour of the voices classified as non-interrogative. Kernels were then normalized by dividing them by the absolute sum of their values and then averaged over all participants for visualization. A one-way repeated-measures ANOVA was conducted on the temporal kernels to test for an effect of segment on pitch shift, and posthocs were computed using Bonferroni-corrected Tukey tests.

**Fig 4.**
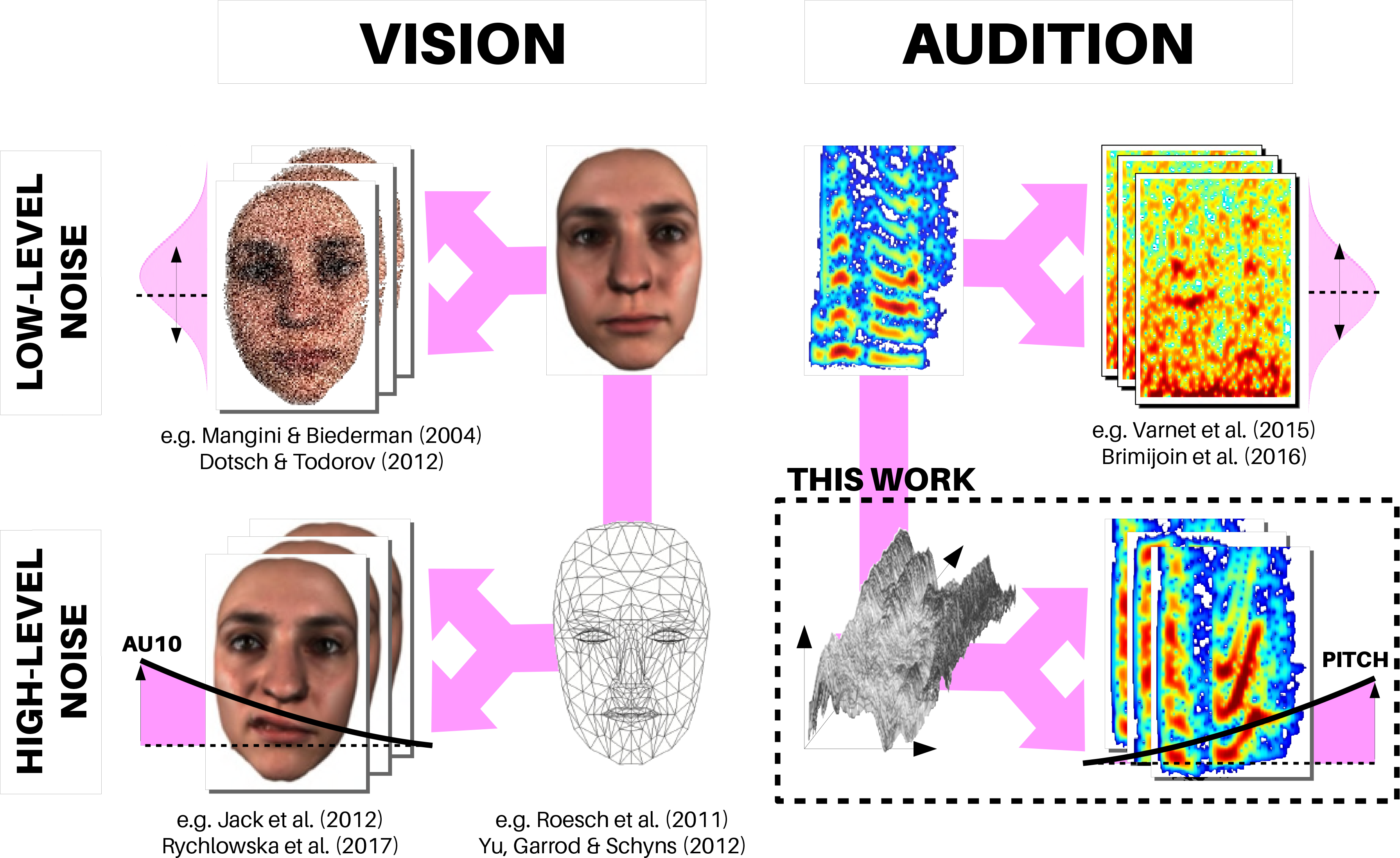
Mental representation of interrogative prosody. Left: Average first-order kernel derived by reverse-correlation on N=5 participants, showing an increase of pitch at the end of the second syllable. Right: individual kernel from each of the participants.

### Results

Observers’ responses revealed mental representations of interrogative prosody showing a consistent increase of pitch at the end of the second syllable of the stimulus word (Fig. 4-left), as reflected by a main effect of segment index : F(6,24)=35.84, p=7.8e-11. Pitch shift at segment 5 (355ms) was significantly different from all other segment locations (all ps <0.001). The pattern was remarkably consistent among participants, although all participants were tested on separate set of random stimuli (Fig. 4-right).

## Case-study (2): musical rhythm

To illustrate another use of CLEESE, this time with musical stimuli and the time-stretching functionality, we describe here a second reverse-correlation experiment about the perception of musical rhythm and expressive timing.

The study of how participants perceive or accurately reproduce the rhythm of musical phrases has informed such domain-general questions as how humans represent sequences of events [46], internally measure speed and tempo [47] and entrain to low- and high-frequency event trains [48]. For instance, it is often observed that musicians tend to lengthen notes at the ends of musical phrases [49] and that even non-musicians anticipate such changes when they perceive music [50]. However, the cognitive structures that govern a participant’s representation of the rhythm of a given musical passage are difficult to uncover with experimental methods. In [50], it was accessed indirectly by measuring the ability to detect timing errors inserted at different positions in a phrase; in [51], participants were asked to manually advance through a sequence of musical chords with a key press, and the time dwelt on each successive chord was used to quantify how fast they internally represented the corresponding musical time. Here, we give a proof of concept of how to use CLEESE in a reverse-correlation experiment to uncover what temporal contour drives participants’ judgement of a rhythmically competent/accurate rendition of the well-known song *Happy birthday*.

## Methods

### Stimuli

One female singer recorded a *a capella* rendering of the first phrase of the French folk song “Joyeux anniversaire” (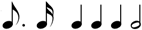; translation of English song \Happy birthday to you” [52]). We used CLEESE to artificially manipulate the timing of the recording by stretching it between different breakpoints. First, we manually identified the time onset of each sung note in the phrase, and use these positions as breakpoints. Second, the original temporal contour was artificially flattened (i.e. all notes were stretched/compressed to have identical duration ♩♩♩♩♩♩), while preserving the original pitch of each note. The duration of this final stimulus was 3203ms. Then, we added Gaussian “temporal noise” (i.e. time-stretching) to each note by sampling stretch values at 6 successive time-points, using a normal distribution centered at 1 (SD = 0.4; clipped at ± 1.6 SD), using a square BPF with a transition time of 0.1 s. The resulting stimuli were therefore sung variants of the same melody, with the original pitch class, but random rhythm (and overall duration).

### Procedure

Pairs of these randomly-manipulated sung phrases were presented to N=12 observers (male: 6, M=22yo), all native French speakers with self-reported normal hearing. Five participants had previous musical training (more than 12 years of instrumental practice) and were therefore considered as musicians, the other seven participants had no musical training and were considered as non-musicians. Participants listened to a pair of two randomly-modulated phrases and were asked which of the two versions was best sung/performed. Inter-stimulus interval in each trial was 500 ms, and inter-trial interval was 1s. Each participant was presented with a total of 313 trials. There were 280 different trials and the last block of 33 trials was repeated twice (in the same order) to estimate the percentage of agreement. After the test, all participants complete the rhythm, melody, rythm-melody subtests of the Profile of Music Perception Skills (PROMS, [53]), in order to quantify their melodic and rhythmic perceptual abilities.

### Apparatus

Same as previous section.

### Analysis

A first-order temporal kernel [45] (i.e., a 6-points vector) was computed for each participant, as the mean stretch contour of the phrases classified as ‘good performances’ minus the mean stretch contour of the phrases classified as ‘bad performances’^6^.

Kernels were then normalized by dividing them by the absolute sum of their values and then averaged over all participants. A one-way repeated-measures ANOVA was conducted on the temporal kernels to test for an effect of segment on time-stretch, and posthocs were computed using Bonferroni-corrected Tukey tests. In addition, the amount of internal noise for each subject was computed from the double-pass percentage of agreement, using a signal detection theory model including response bias and late additive noise [54, 55].

## Results

**Fig 5.**
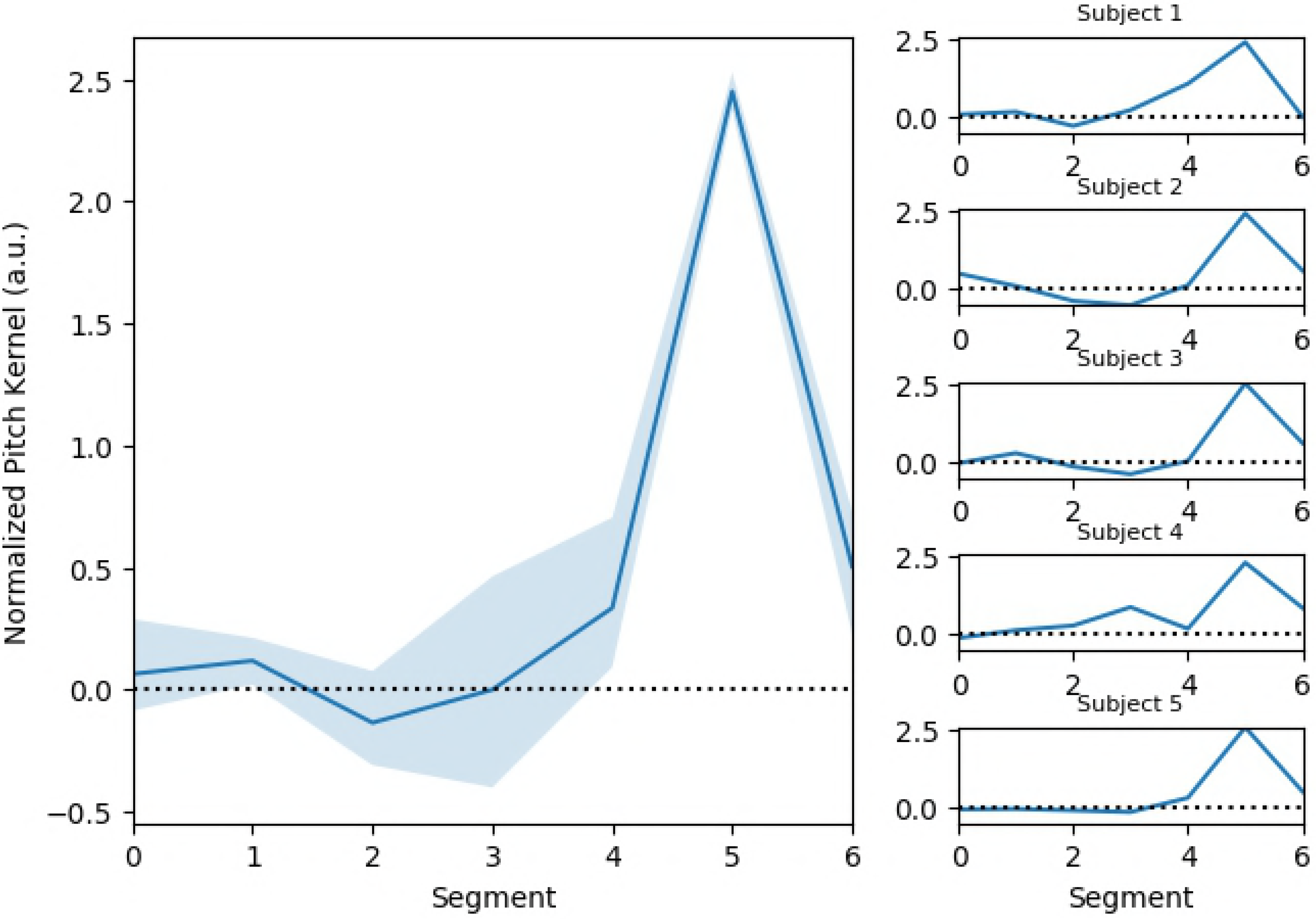
Mental representation of the rhythm of a well-known song. Left: Average first-order kernel derived by reverse-correlation on N=12 participants, superimposed with the theoretical pattern (dotted line), showing how the internally represented pattern departed from the theoretical one. Right: log-values of internal noise estimated from the double-pass technique are negatively correlated with the participant’s degree of musical skills (average score obtained for the 3 sub-tests of PROMS); r=−0.65, p=0.02.

Observers’ responses revealed mental representations of note durations that significantly evolve through time (Fig. 5-left) with a main effect of time index : F(5,40)=30.7, p=1e-12. The ideal theoretical timing contour (0.75-0.25-1-1-1-2), as given by the score of this musical phrase, was converted in time-stretch units by taking the log of these values and was superimposed on Fig. 5. Several deviations from this theoretical contour are worth noting. First, both musicians and non-musicians had shortened representations for the first note of the melody 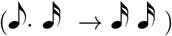. Second, in an unpredicted effect, the analysis revealed a significant interaction of time index and participant musicianship: F(5,40)=2.5, p=.045. The mental representations of a well-executed song for the non-musician participants had longer durations on the third note (“happy BIRTH-day to you”) compared with musicians participants. Needless to say, this difference between musicians and non-musicians is only provided for illustrative purposes, because of the small-powered nature of this case-study, and its experimental confirmation and interpretation remain the object of future work. Finally, both musicians and non-musicians had similarly shortened representations for the duration of the last note 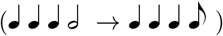, a pattern which may appear at odds with the phrase-final lengthening usually seen in both musicians and non-musicians [49, 51]. It is possible that smaller time-stretch values for this time point result from the participants’ overestimating the duration of the final note, an aspect that remains to be tested in future studies.

One should note that it is complicated to quantitatively compare the time-stretch contours obtained with reverse-correlation with an ideal theoretical pattern. First, time-stretch factors derived by reverse-correlation have arbitrary scale, and the ideal pattern is therefore be represented at an arbitrary position on the y-axis, and rescaled to have the same (max-min) stretch range as the one obtained experimentally. Second, because participants’ task is to infer about the best overall rhythm and not individual note duration, the time-stretch values at individual time points are not perceptually independent (e.g., if the first two notes of a stimulus are long, participants may infer a slower beat and expect even longer values for the following notes). For these reasons, the ideal rhythmic pattern superimposed on these measured patterns should simply be taken as an illustration of how note durations theoretically evolve from one note (i) to the next (i+1). Further theoretical work will be needed to best analyze and interpret duration kernels derived from such reverse-correlation experiments.

In addition, internal noise values computed from the repeated block of this experiment indicate that subjects did not behave at random but rather relied on a somewhat precise mental representation of the ideal temporal contour (mean internal noise of 1.1 (SD=1.1) in units of external noise; comparable with the average value of 1.3 obtained in typical low-level sensory psychophysical detection or discrimination tasks, see [55]). In an exploratory manner, we asked whether these internal noise values would correlate with the participant’s degree of musical skills (average score obtained for the 3 sub-tests of PROMS). We found a significant negative correlation (r=-0.65, p=0.02), suggesting that low musical skills are associated with a more variable mental/memory representation of the temporal contour of this melody 5-right). Because this correlation may also be driven by the amount of attention that participants had both in the PROMS and in the reverse-correlation tasks (which may be similar), further work is required to determine the exact nature of this observed relationship.

## Discussion

By providing the ability to manipulate speech and musical dimensions such as pitch, duration and amplitude in a parametric, independent manner in the common environment of the Python programming language, the open-source toolbox CLEESE brings the power of data-driven / reverse-correlation methods to the vast domain of speech and music cognition. In two illustrative case-studies, we have used CLEESE to infer listeners’ mental representations of interrogative intonation (rising pitch at the end of the utterance) and of the rhythmic structure of a well-known musical melody, and shown that the methodology had potential to uncover potential individual differences linked, e.g., to participant’s training or perceptual abilities. As such, the toolbox and the associated methodology open avenues of research in communicative behavior and social cognition. As a first application of CLEESE, we have recently used a reverse-correlation paradigm to uncover what mental representations of pitch profiles underlie judgements of speaker dominance and trustworthiness in short utterances like the word ‘hello’ [31]. The technique allowed to establish that both constructs corresponded to robust and distinguishing pitch trajectories, which remained remarkably stable whether males or female listeners judged male or female speakers. Other potential questions include, in speech, studies of expressive intonation along all characteristics of pitch, speed and amplitude, judgements of emotions (e.g. being happy, angry or sad) or attitudes (e.g. being critical, impressed or ironic); in music, studies of melodic and rythmic representations in naive and expert listeners, and how these may differ with training or exposure. Beyond speech and music, CLEESE can also be used to transform an study non-verbal vocalizations, such a infant cries or animal calls.

By measuring how any given individual’s or population’s mental representations may differ from the generic code, data-driven paradigms have been especially important in studying individual or cultural differences in face [2, 56] or lexical processing [23]. By providing a similar paradigm to map mental representations in the vast domain of speech prosody, the present technique opens avenues to explore e.g. dysprosody and social-cognitive deficits in autism-spectrum disorder [57], schizophrenia [58] or congenital amusia [59], as well as cultural differences in social and affective prosody [60].

Because CLEESE allows to create random variations among the different dimensions of both speech and musical stimuli (pitch, time, level), it also opens possibilities to measure the amount of internal noise (e.g. using a double-pass technique as in the case-study 2 here) associated to the processing of these dimensions in many various high-level cognitive tasks. This is particularly interesting because it provides a quantitative way to (1) demonstrate that participants are not doing the task at random (which is always an issue in this type of high-level task where there is no good/bad answers that would lead to an associated d-prime value) and (2) investigate which perceptual dimensions are cognitively processed with what amount of noise.

The current implementation of CLEESE, and its application to data-driven experiments in the auditory modality, leaves several methodological questions open. First, in the current form, the temporal dynamics of the noise/perturbations is purely random, and changes of pitch/time or amplitude are independent across segments (a classical assumption of the reverse-correlation technique [4]). This assumption may create prosodic patterns that are not necessary realizable by the human voice and thus bias the participant responses for those trials. In vision, recent studies have manipulated facial action units with a restricted family of temporal profiles parametrized in acceleration, amplitude and relative length [25]. In a similar manner, future versions of CLEESE could constrain prosodic patterns to correspond more closely to the dynamics of the human voice or to the underlying production process, e.g. not aligned with arbitrary segment locations but with the underlying phonemic (or musical) structure.

Another possible direction for improvement is the addition of other high-level manipulation dimensions. As an example, spectral envelope manipulation (optionally formant-driven) allows powerful transformations related to timbre, speaker identity and even gender which all could be manipulated by CLEESE [61]. At an even higher abstraction level, research could consider more semantically-related features (such as the mid-level audio descriptors typically used in machine learning methods) or even feature-learning approaches that would automatically derive the relevant dimensions prior to randomization [62]. Adding these new dimensions will likely require a trade-off between their effectiveness for stimuli randomization and their suitability for physical interpretation.

Finally, on a purely technical level, improvements of the current phase vocoder implementation in CLEESE may include additional modules such as envelope or transient preservation to further improve the realism of the transformations.

## Supporting information

All experimental data and analysis scripts (in the form of Jupyter notebook) for the two case-study experiments are made available as supplementary material

## Acknowledgments

Joint first authors JJB and EP made equal contributions. The authors thank Etienne Thoret, Leo Varnet, Vincent Isnard for useful discussions on the audio signal processing aspects of this work, as well as Pascal Belin, Frederic Gosselin, Philippe Schyns and Rachael Jack for discussions on data analysis. We thank Louise Vasa for help with data collection. All data collected at the Sorbonne-Université INSEAD Center for Behavioural Sciences. This study was supported by ERC Grant StG 335536 CREAM to JJA.

## Footnotes

1 named after British actor John Cleese, with reference to the “Ministry of Silly Talks”

2 Available upon free registration at http://forumnet.ircam.fr/product/cleese

3 Note that CLEESE can also be used to create spectral transformations, a technique not described here - see [32]

4 In more details, CLEESE implements variants of these procedures: time-stretching is implemented by taking a time-varying hop size at the analysis stage before the STFT and a constant hop size at the synthesis stage, and pitch-shifting is implemented as time stretching followed by resampling

5 we created these “high” and “low” pitch categories to facilitate participants’ task, but they are not necessary

6 Because stretch factors are ratio, the kernel was in fact computed in the log stretch domain, as *mean* log *t*^+^ − *mean* log *t*^−^, where *t*^+^ and *t*^−^ are the contours of selected and non-selected trials, resp.

